# A Multidomain Lens on the Temporal Dynamics of Surface Microbial Communities in the Southern Ocean (2013–2019)

**DOI:** 10.1101/2025.03.27.645779

**Authors:** Laiza C. Faria, Yubin Raut, Jesse McNichol, Nathan L. R. Williams, Jed A. Fuhrman, Camila N. Signori

**Affiliations:** Department of Biological Oceanography, Oceanographic Institute, University of São Paulo, São Paulo, Brazil; Department of Earth, Atmospheric, and Planetary Sciences, Massachusetts Institute of Technology, Cambridge, MA, USA; Department of Biology, St. Francis Xavier University, Antigonish, NS, Canada; Department of Biological Sciences, University of Southern California, Los Angeles CA, USA

**Keywords:** Microbial Oceanography, Metabarcoding, Antarctic Peninsula, Prokaryotic stability, Eukaryotic variability

## Abstract

Marine microorganisms are vital to biogeochemical cycles and food web dynamics, with their community structure shaped by environmental factors such as temperature, light, and salinity. While microbial dynamics in the western Antarctic Peninsula are relatively well-studied, the northwestern region remains underexplored, particularly in long-term, multidomain analyses. To fill this gap, we investigated microbial communities encompassing all three domains of life (Bacteria, Archaea, and Eukarya) in the Northwestern Antarctic Peninsula. Using the universal primer set 515Y/926R, we sequenced unfractionated seawater from ten sites over a six-year period (2013–2019). Environmental parameters, temperature and salinity, showed minimal variation across the study. However, microbial diversity and composition, especially among eukaryotic phytoplankton, displayed significant temporal changes among seasons and years. The prokaryotic community, by contrast, was relatively stable, with *Gammaproteobacteria*— particularly the *Nitrincolaceae* family—maintaining high relative abundance throughout all sampling periods, but a few distinct ASVs. In contrast, no eukaryotic group exhibited consistently high relative abundance across sampling periods. The summer of 2016, marked by a strong El Niño event, presented the most distinct microbial community structure, underscoring the sensitivity of these communities to extreme climatic conditions. These results highlight the importance of integrated, long-term studies to better understand the dynamics, interactions, and resilience of microbial ecosystems in the rapidly changing Antarctic environment.

**IMPORTANCE:** This study provides a unique long-term perspective on microbial community dynamics in the Northwestern Antarctic Peninsula, a region still poorly explored through a multidomain lens. By investigating the temporal variability of Bacteria, Archaea, and Eukaryotes over six years, we reveal distinct stability patterns between these groups, with phytoplankton showing the highest variability and prokaryotes remaining relatively stable. The strong response of the microbial community to the 2016 El Niño event highlights its sensitivity to extreme climate conditions, reinforcing the importance of understanding how Antarctic ecosystems will respond to future climate shifts. The consistent presence of *Nitrincolaceae*, a key bacterial taxon, suggests its ecological relevance in the region, while fluctuations in phytoplankton composition may impact food web dynamics. These findings emphasize the need for continued long-term monitoring to predict how microbial communities will adapt to environmental changes, which is crucial for assessing the future functioning of polar marine ecosystems.

## INTRODUCTION

Microorganisms play crucial roles in marine ecosystems, driving biogeochemical cycles and forming the foundation of marine food webs, which are vital for sustaining life in the oceans (Amaral-Zettler et al., 2010; Azam et al., 1983; Sunagawa et al., 2015). They play a key role in cycling essential elements such as carbon, nitrogen, oxygen, sulfur, and iron. Within the carbon cycle, their functional processes include photosynthesis, methanogenesis, respiration, and fermentation (Falkowski et al., 2008). In marine food webs, autotrophic microorganisms—both photosynthetic and chemosynthetic—constitute the base of the trophic structure. Here, we define the microbial community as encompassing marine plankton from all three domains of life: Bacteria, Archaea, and Eukarya.

The Northwest Antarctic Peninsula (NAP) is a highly dynamic region where seasonal changes in solar radiation and sea ice dynamics drive strong fluctuations in primary productivity (Venables et al., 2013). Phytoplankton blooms occur in response to variations in sea ice extent and water column stability (Schofield et al., 2018). However, the NAP is also experiencing rapid environmental changes due to climate change, affecting the atmosphere, cryosphere, and ocean (Stammerjohn et al., 2008). Large-scale climate patterns such as the El Niño-Southern Oscillation (ENSO) and the Southern Annular Mode (SAM) further influence the region’s biogeochemistry and ecosystem dynamics(Stammerjohn et al., 2008). These ongoing changes make the NAP an ideal setting for studying the impact of climate change on polar marine systems (Henley et al., 2019).

Marine microorganisms are subject to a range of biotic and abiotic factors that vary across temporal and spatial scales, driving changes in Plankton community composition (Fuhrman, 2009; Fuhrman et al., 2015). Temporal variations in these communities can occur over daily, monthly, or yearly timescales. In the surface layer of seawater, this phenomenon occurs primarily due to natural variations in temperature and sunlight hours, which often trigger responses in the phytoplankton community, including cyanobacteria and eukaryotic photosynthetic organisms. In contrast, the prokaryotes, consisting of non-autotrophic bacteria and archaea remineralizers, are often more stable across time compared to the eukaryotic fraction (Caporaso et al., 2012; Y.-C. Yeh & Fuhrman, 2022).

Antarctic microbial communities are strongly influenced by depth-dependent environmental fluctuations, including sunlight, temperature, and nutrient availability. In the oceanic region northwest of the Antarctic Peninsula, seasonal variations in solar radiation, luminosity, and temperature play a crucial role in shaping the structure of surface microbial communities (Ducklow et al., 2013; Smetacek & Nicol, 2005). These communities, which are more exposed to short-term atmospheric changes such as sunlight exposure, temperature shifts, and wind, are dominated by taxa such as *Alteromonadales*, *Rhodobacterales*, *Flavobacterales*, haptophytes, and diatoms (Luria et al., 2014; Ozturk et al., 2022; Signori et al., 2018). In contrast, deep-water microbial assemblages are primarily structured by water masses with longer residence times (Ghiglione et al., 2012).

Members of the marine microbial community also interact metabolically, complementing each other by exchanging metabolic intermediates and cellular building blocks to achieve collective metabolism and consequently, playing crucial roles in nutrient cycling (Huelsmann et al., 2024). In the Antarctic region, interactions between marine eukaryotic and prokaryotic taxa have been shown to influence each other’s relative abundances, as demonstrated through both in situ studies and experimental research (Dawson et al., 2023; Delmont et al., 2014). Adopting a multidomain perspective provides a more integrated understanding of these interactions and the overall dynamics of plankton communities, shedding light on the broader impacts of environmental changes.

Some studies have investigated the short-term temporal variation of microbial communities in the southern part of the western side of the maritime Antarctic Peninsula (Ducklow et al., 2013; Grzymski et al., 2012; Luria et al., 2016). However, there is still a significant gap in research focusing on the northwestern part of the maritime Antarctic Peninsula. More importantly, long-term time series studies, spanning several years, on how these communities respond to environmental changes remain scarce. This type of research is particularly relevant in the context of the rapid environmental changes this region has been experiencing (Clarke et al., 2007; Ducklow et al., 2013). Lastly, these studies have only considered either the prokaryotic or eukaryotic components of the microbial community, without assessing the entire community under a single denominator.

Therefore, we aimed to characterize the microbial community from all three domains of life from the northwestern side of the Antarctic Peninsula to reveal how different plankton groups, both individually and collectively, respond to environmental conditions such as temperature and salinity and fluctuate over time. To do this, we sequenced unfractionated seawater from ten different sites in the Northwest Antarctic Peninsula (NAP) region with a universal primer set (515Y/926R) (Parada et al., 2016) during a 6-year time series from 2013 to 2019. By using a universal primer set (515Y/926R) (Parada et al., 2016), we were able to assess the dynamics of the Bacteria, Archaea, and Eukarya under a common framework. This approach provided valuable insights into the complex interactions within the microbiome using a multidomain perspective and its resilience to environmental changes.

## RESULTS

### ENVIROMENTAL PARAMETERS

Salinity values remained relatively stable over time and across areas, averaging around 34 PSU. Excluding an outlier (near 30 PSU) in Spring 2013, samples taken in Late Summer 2014 exhibited the lowest salinity values, nearing 32 PSU, while Spring 2013 had the highest, with values approaching 35 PSU (see Figure 1a). Temperature ranged from -1.77°C to 2.48°C across all sampling periods. The lowest temperatures were recorded during Spring 2013 and 2014, the only two periods when all values were below 0°C. Temperatures above 1°C were observed in Summer and Late Summer 2015, Summer 2017, and Summer 2019, with Summer 2017 being the warmest period, also showing the greatest variation between areas (see Figure 1b). All data are available in Supplementary Table 1.

**Figure 1:**
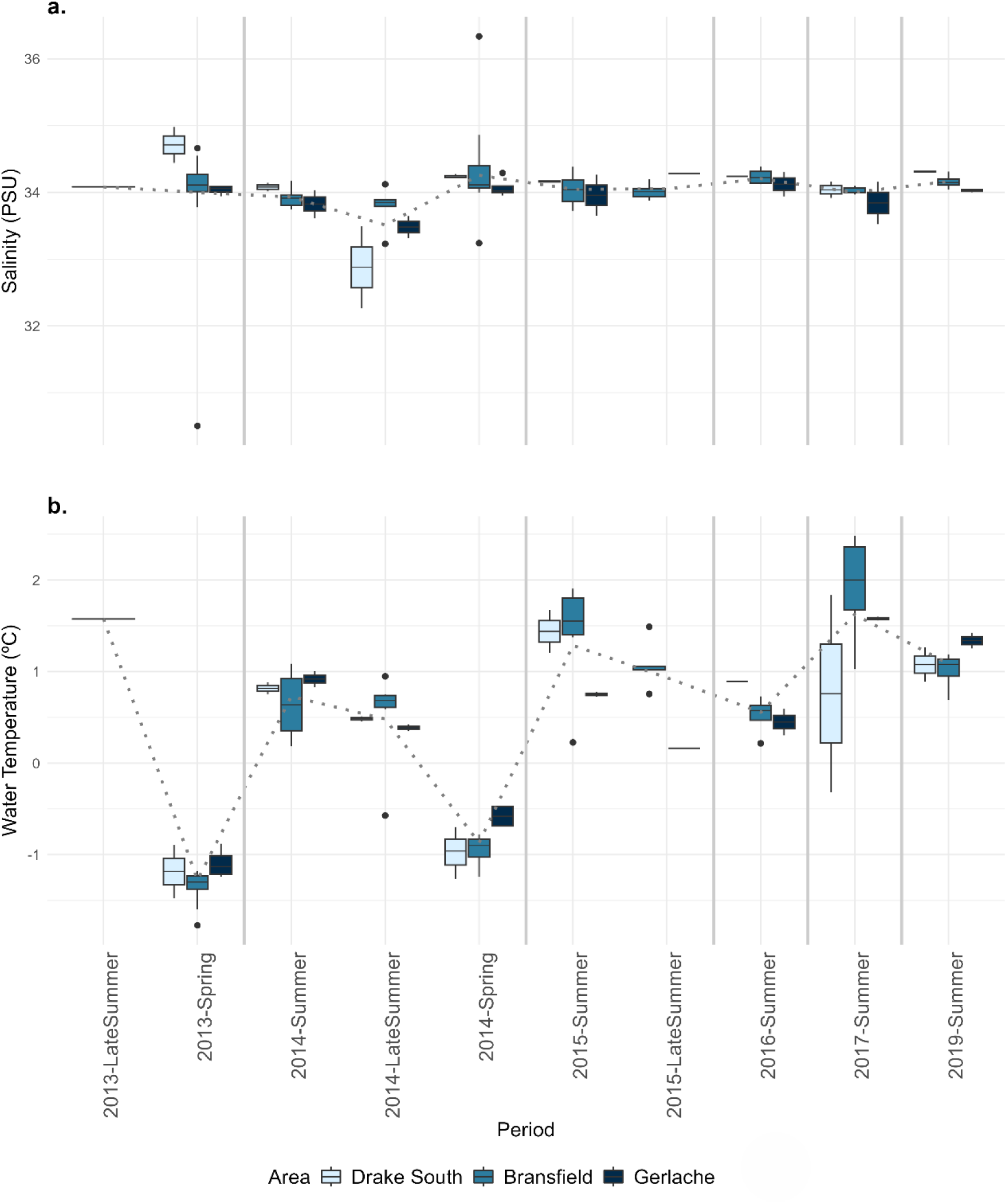
Boxplots showing the variation in physicochemical properties. a. Salinity (PSU) and b. Temperature (°C), across all sampling periods (x-axis). Samples are grouped by sampling area: the lightest blue represents Drake South, medium blue represents Bransfield Strait, and the darkest blue represents Gerlache Strait. Solid lines separate the years, and the dashed line shows the variation of the average across all periods.

### PLANKTON COMMUNITY ALPHA AND BETA DIVERSITY

The Shannon H-Index revealed distinct patterns of alpha diversity variation across different microbial groups and the total community over the sampling periods and areas (Figure 2). Among all groups, the photosynthetic eukaryotic community (Chloroplast 16S) showed the highest variability, with Shannon index values fluctuating by more than 700%—representing the relative difference between the lowest and highest values observed across different periods—between the least diverse period (Summer 2016) and the most diverse (Spring 2014). In particular, during Summer 2017 in Bransfield, the variation peaked at 600% (see Figure 2). Similarly, the total eukaryotic community, encompassing both photosynthetic and non-photosynthetic organisms, displayed substantial diversity fluctuations, with a maximum variation of 660% between periods and areas. In contrast, the prokaryotic community demonstrated the highest stability in alpha diversity across sampling areas and periods, with a maximum variability of only 50%, as calculated at the ASV level. When combining prokaryotic and eukaryotic communities (photosynthetic and non-photosynthetic), the total microbial community (MultiDomain) also exhibited relative stability, with a variability of approximately 52% (see Figure 2).

**Figure 2.**
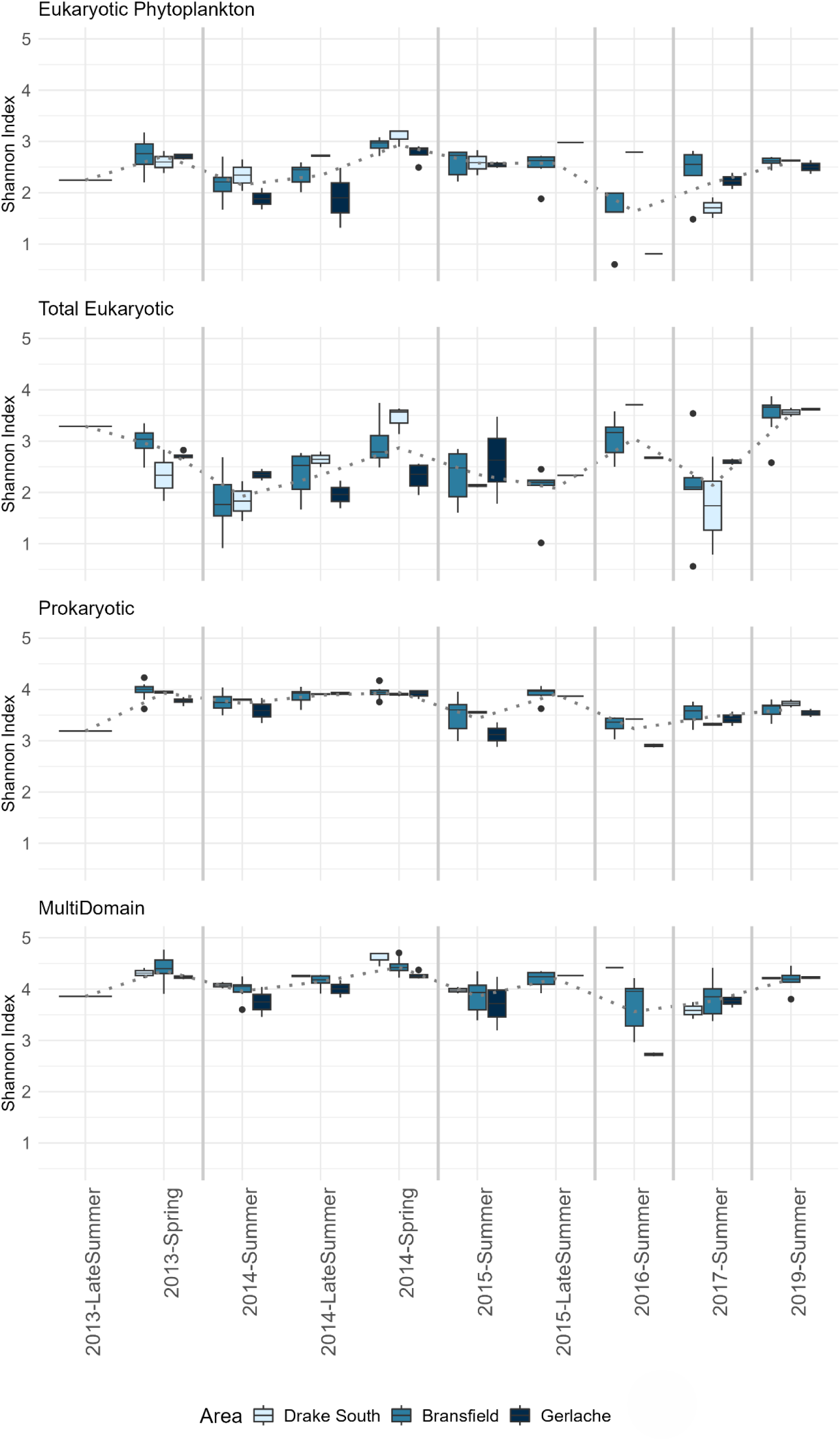
Boxplots of Shannon diversity index values for microbial communities across different periods and areas in the Southern Ocean. The panels represent diversity indices for Eukaryotic Phytoplankton, Total Eukaryotic, Prokaryotic, and MultiDomain. Samples are grouped by sampling area: the lightest blue represents Drake South, medium blue represents Bransfield Strait, and the darkest blue represents Gerlache Strait. Solid lines separate the years, and the dashed line shows the variation of the average across all periods.

A temporal analysis revealed a trend of higher diversity during spring seasons (2014 and 2015) and lower diversity during the summer seasons. Within each period, most groups showed consistent patterns of variation, either increasing or decreasing in diversity together. Notably, during the Summer 2016 period, a marked decrease in diversity was observed in the photosynthetic eukaryotic community, the prokaryotic community, and the total microbial community, with the average diversity being lower compared to other periods. However, an opposing trend was noted in the total eukaryotic community (including both photosynthetic and non-photosynthetic organisms), which experienced an increase in diversity during the same period (see Figure 2).

The composition of each plankton community varied across both spatial and temporal scales. Among Eukaryotic Phytoplankton, the 2016 community showed the greatest dissimilarity compared to other sampled years (Figure 3). Colder samples tended to cluster together, although their dissimilarity from warmer samples was not statistically significant. The Total Eukaryotic community exhibited the highest dissimilarity among samples collected within the same year. Similar to the Eukaryotic Phytoplankton, colder samples clustered together but were not significantly different from warmer ones. Notably, the Total Eukaryotic community maintained at least 50% similarity between samples from 2016 and 2019.

**Figure 3.**
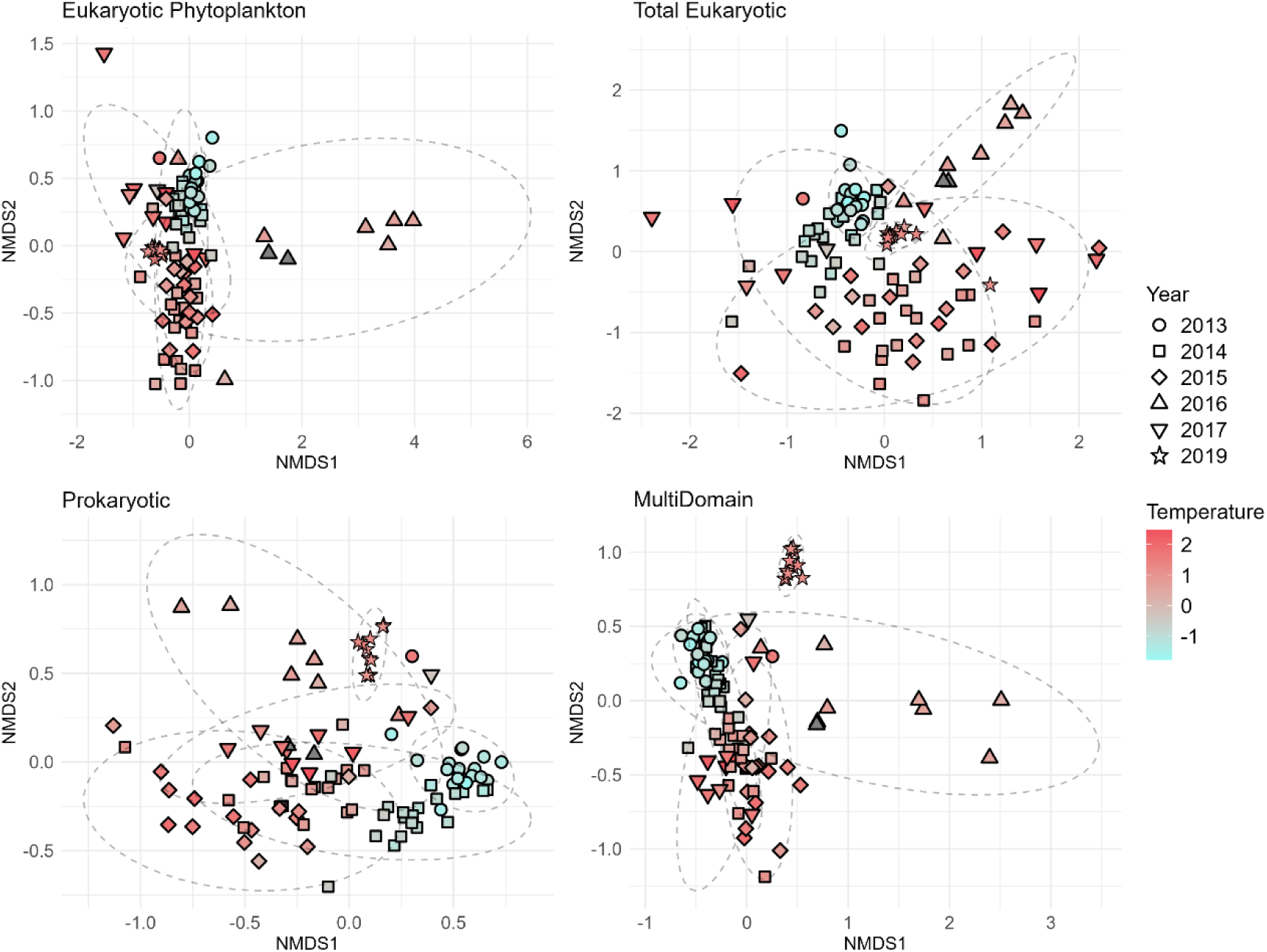
Nonmetric multidimensional scaling (nMDS) ordination based on Bray–Curtis distances for Eukaryotic Phytoplankton, Total Eukaryotic, Prokaryotic groups, and the MultiDomain community. Shapes represent different years, while temperature is displayed as a gradient, with cooler temperatures in blue and warmer temperatures in red. Ellipses indicate groups of samples sharing at least 50% similarity.

The Prokaryotic community was the most internally consistent, with temperature creating a clearer separation among samples. When analyzing the entire community together, the MultiDomain plot revealed a distribution pattern similar to that of the Eukaryotic Phytoplankton community. However, in the MultiDomain analysis, both 2016 and 2019 stood out for their dissimilarity. While 2016 samples were more internally variable, 2019 samples were highly similar to each other but distinct from other years.

The PERMANOVA test results revealed that the sampling period had a highly significant effect across all community fractions, accounting for 52% of the variation in Eukaryotic Phytoplankton, 40% in the Total Eukaryotic community, 56% in Prokaryotes, and 57% in the MultiDomain ASV composition. Additionally, temperature, area, and the interaction between period and area were significant factors (p < 0.05), though their effects were less pronounced (Table 1). In contrast salinity was not significant (p > 0.05).

**Table 1:** R^2^ values for sources of overall variation determined by the PERMANOVA test for the Eukaryotic Phytoplankton, Total Eukaryotic, Prokaryotic groups, and the MultiDomain community. Only results with p-values < 0.05 are included.

|  | **Eukaryotic**<br>**Phytoplankton** | **Total**<br>**Eukaryotic** | **Prokaryotic** | **MultiDomain** |
| --- | --- | --- | --- | --- |
| Period | 0.52 | 0.40 | 0.56 | 0.57 |
| Temperature | 0.03 | 0.02 | 0.01 | 0.02 |
| Area | 0.02 | 0.03 | 0.03 | 0.03 |
| Period:Area | 0.11 | 0.14 | 0.10 | 0.10 |

### MICROBIAL COMMUNITY COMPOSITION

As presented above, the prokaryotic community demonstrated the highest stability among microbial groups. *Gammaproteobacteria*, particularly the *Nitrincolaceae* (formerly *Oceanospirillaceae*) family, were consistently present across all sampling periods, maintaining a mean relative abundance exceeding 2%. Similarly, *Alphaproteobacteria* and *Bacteroidia* exhibited mean relative abundances above 2% in most sampled periods, with exceptions for *Alphaproteobacteria* in Spring 2014 and Summer 2019, and for *Bacteroidia* in Spring 2013, Summer 2016, and Summer 2019 (Figure 4).

**Figure 4.**
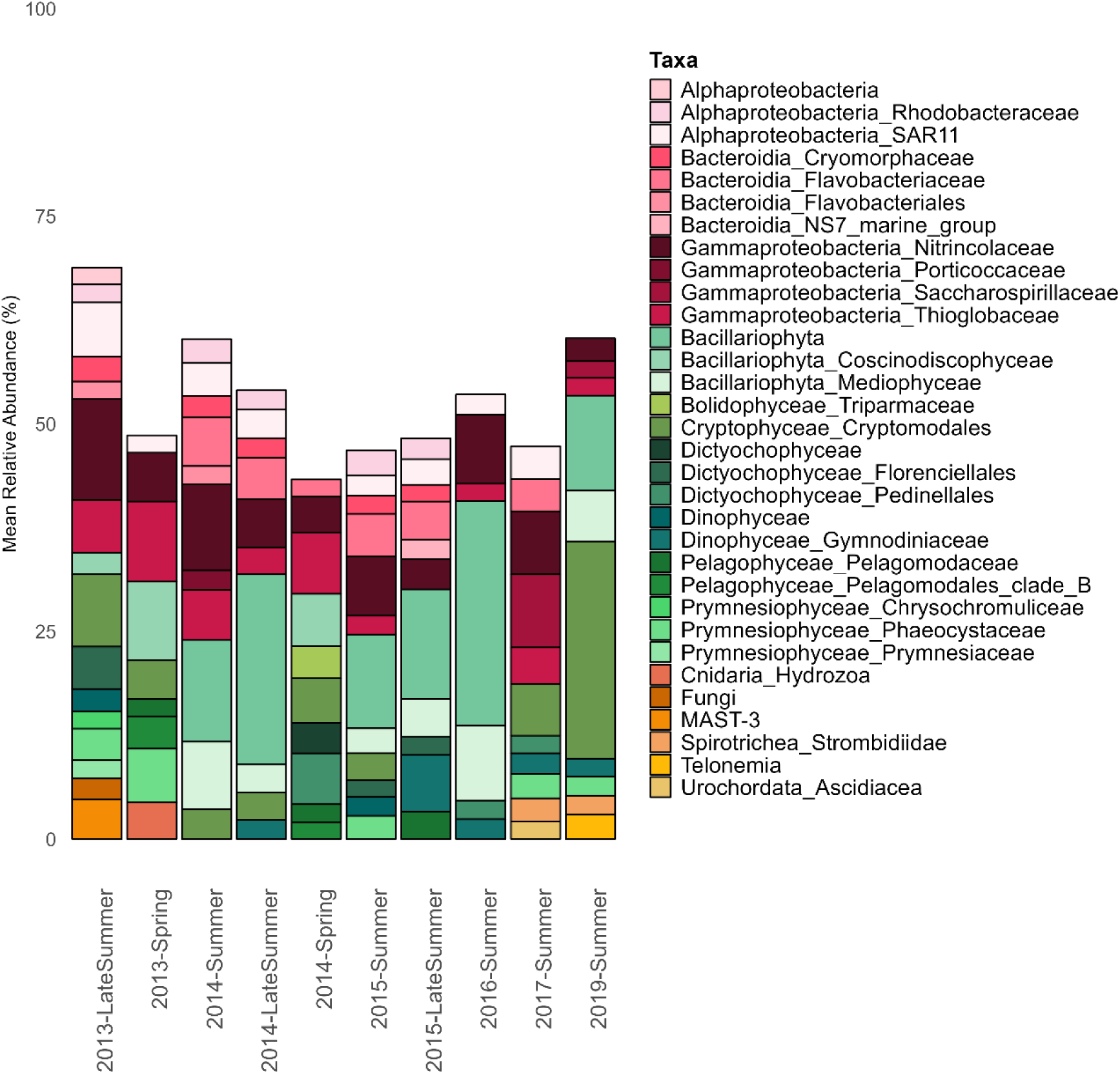
Mean relative abundance of the most abundant taxa (average relative abundance exceeding 2% per period). Bacterial groups are shown in shades of pink, phytoplankton groups in shades of green, and other eukaryotes in shades of orange.

In contrast to this stability, the eukaryotic phytoplankton community showed considerable variation. Many groups appeared across different sampling periods but lacked consistency over time. Remarkably, no eukaryotic group (both photosynthetic and non-photosynthetic), not even at the order level, was consistently present throughout all sampled periods.

Periods such as Late Summer 2013, Summer and Late Summer 2014, Summer 2016, and Summer 2019 were characterized by the most abundant taxa (the ones with mean relative abundance higher than 2%) accounting for more than 50% of the community composition. Spring 2013 and 2014 exhibited the highest proportions of bacteria, whereas Summer 2016 and 2019 showed a greater dominance of phytoplankton. Notably, some periods were dominated by phytoplankton groups; Late Summer 2014 and Summer 2016 by *Bacillariophyta* (22.9% and 27.1%, respectively), and Summer 2019 by *Cryptophyceae Cryptomodales* (26.1%), respectively, likely indicating phytoplankton blooms during the sampling period (Figure 4).

The core microbiome was defined at the Family level as all families present in all samples from all periods and was composed of 14 microbial groups *Flavobacteriales; Cryomorphaceae; Cryptomonadales; Flavobacteriaceae; Florenciellales; Magnetospiraceae; Nitrincolaceae; Phaeocystaceae; Porticoccaceae; Rhodobacteraceae; Rickettsiales; SAR11; SAR86_clade* and *Thioglobaceae*. Some groups, such as *SAR11* and *Nitrincolaceae*, exhibited lower variability and higher relative abundances, whereas *Thioglobaceae* and *Cryptomonadales*, despite being present in all samples across all periods, showed variations in their mean relative abundance. *SAR86_clade*, *Rickettsiales*, and *Magnetospiraceae*, although consistently present, maintained low relative abundances throughout all periods.

Network analyses based on significant Spearman correlations greater than 0.6 or less than -0.6 revealed predominantly positive associations both within and between domains (Figure 5). Prokaryotes exhibited the highest number of positive interactions overall, with *Chloroflexi* standing out as the group with the most positive associations above the predefined threshold, totaling 23 positive interactions with other groups, followed by *Acantharea* with 21 positive interactions and *Defluviicoccales* with 20 positive interactions. Notably, the only negative interactions observed occurred within domains, specifically prokaryotes and eukaryotes. *Flavobacteriaceae* and *Rhodobacterales,* were the two groups with the highest number of negative interactions, with 4 and 3 associations, respectively. On the other hand, the prokaryotic group *Rickettsiales* exhibited exclusively positive interactions, all of which were with photosynthetic eukaryotic groups.

**Figure 5.**
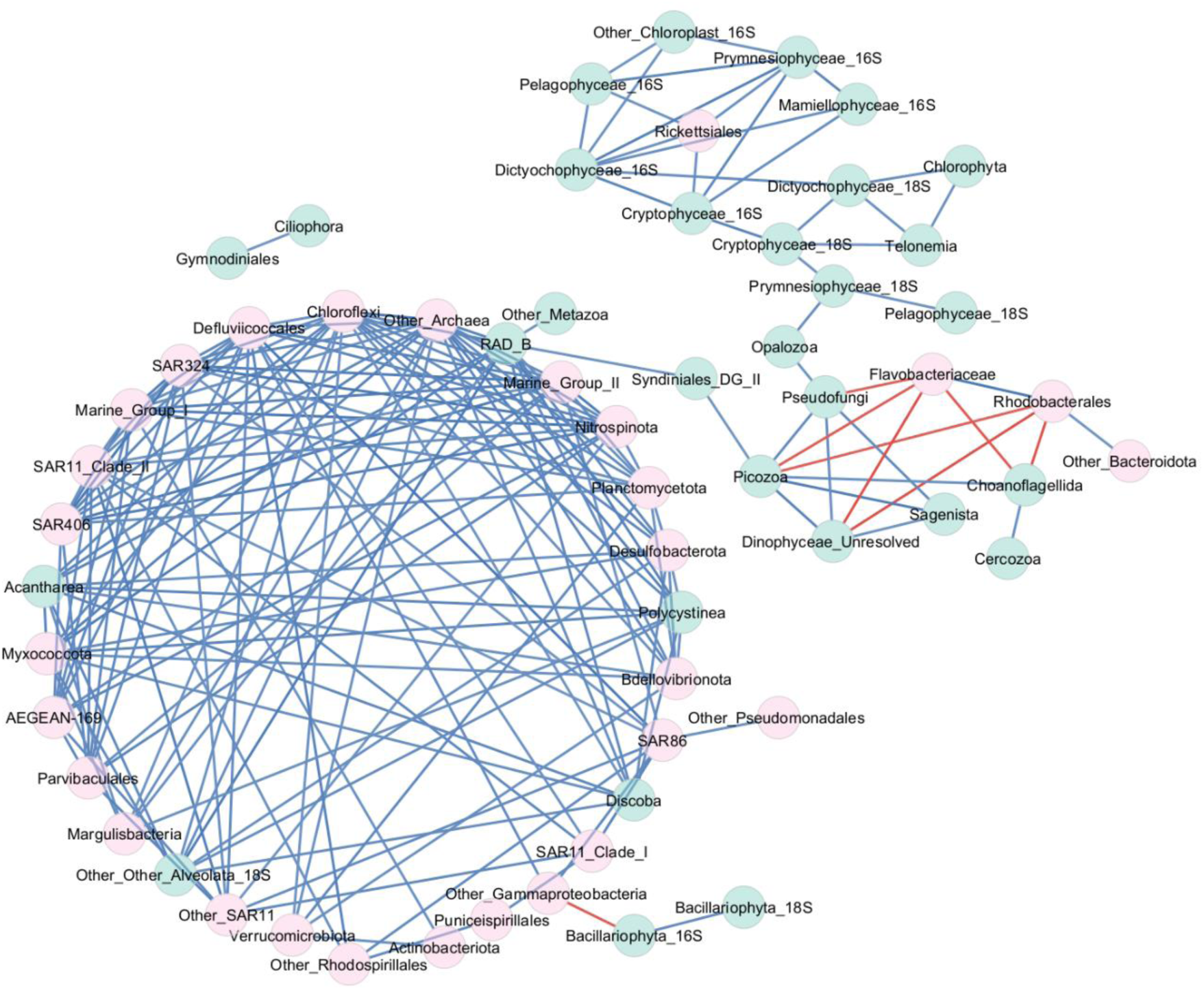
Network analysis where nodes represent ecologically relevant groups (green for eukaryotes and pink for prokaryotes), and edges represent interactions between groups. Blue edges indicate positive interactions, and red edges indicate negative interactions. Only interactions with a p-value < 0.05 and Spearman correlation coefficient > 0.6 or < -0.6 were considered.

## DISCUSSION

The results of this study offer first insights into the temporal dynamics of the surface microbial community using a multidomain lens in the maritime region of the NAP. Our findings demonstrate that different groups of the microbial community—Eukaryotic Phytoplankton, Total Eukaryotic, and Prokaryotic—respond to environmental pressures with varying intensities over time (both seasons and years), with the MultiDomain group representing the combined response of these three groups. Although no significant differences were observed in the environmental parameters analyzed in this study (Figure 1), we detected notable variations in microbial diversity and community composition over time. For all microbial community groups considered in this study, the sampling period explained the largest portion of the variation in community composition, likely reflecting temporal changes in other parameters not analyzed here.

### MICROBIAL COMMUNITY DYNAMICS OVER TIME

After a six-year assessment of surface seawater at 10 sites on the NAP, one of our key findings was that, similar to studies of the microbial community in locations such as Southern California’s San Pedro Ocean Time Series, phytoplankton are the most variable members of the plankton community over time, while prokaryotes remain relatively stable (Y. C. Yeh & Fuhrman, 2022). This pattern holds across different timescales, with the San Pedro Ocean Time Series operating on a monthly scale, whereas our study spans seasonal to annual periods. It also emerges across different spatial scales, as the sampling points in the San Pedro Ocean Time Series are much closer to one another than those in our study.

Higher alpha diversity values (Shannon H-Index) were observed during the two spring sampling periods compared to summer and late summer periods, a trend that was anticipated and has been previously reported in studies conducted in the Northwest and Western Antarctic Peninsula (Figure 2) (Luria et al., 2014, 2016; Signori et al., 2018). This trend is associated with the onset of increasing daylight hours, rising temperatures, and subsequent ice melting, which elevates nutrient concentrations in the water (Luria et al., 2014, 2016). These conditions might promote the growth of groups that were dormant during the winter. However, this trend of higher diversity during spring was not observed in the eukaryotic community, which exhibited its highest values in Summer 2019. This suggests that the eukaryotic community may respond to environmental variations by increasing the abundance of individuals within the same group, possibly as a consequence of higher nutrient availability leading to competitive dominance in a bloom situation.

The summers of 2016 and 2019 were marked by the highest relative abundance of eukaryotes (40.8% and 48.2%, respectively) compared to prokaryotes. While both years were distinctly different from each other and from other periods, the 2019 samples exhibited high within group similarity, in contrast to the variability observed in 2016. Both periods were dominated by specific groups, suggesting the occurrence of blooms: *Bacillariophyta* in Summer 2016 (22.38%) and *Cryptomonadales* in Summer 2019 (26.14%) (Figure 4). *Cryptomonadales* are distinguished by their smaller cell size compared to other phytoplankton groups and their bacterivorous mixotrophic behavior (Deppeler & Davidson, 2017). These traits may provide *Cryptomonadales* with an advantage during periods of lower nutrient availability.

However, Summer 2016 stood out as the most distinct period among those analyzed. During this time, the Eukaryotic Phytoplankton community exhibited the lowest diversity values (mean Shannon H-Index of 1.6) and highest relative abundance (40.8%), whereas the Total Eukaryotic showed a contrasting trend, with increased diversity (mean Shannon H-Index of 3) compared to all other fractions of the community (Figure 2). In this context, the increase in Total Eukaryotic community diversity may indicate a rise in grazer diversity in response to greater food availability. Additionally, the Eukaryotic Phytoplankton and MultiDomain communities displayed the highest levels of dissimilarity (Figure 3). This period coincided with a record-breaking El Niño event, characterized by elevated atmospheric temperatures and pronounced effects on the Southern Ocean (Nicolas et al., 2017; Paolo et al., 2018; Santoso et al., 2017). Notably, a significant diatom bloom was reported in the sampling region during this time, the authors hypothesized that these conditions, driven by higher atmospheric temperatures, accelerated ice melting, leading to increased nutrient concentrations (Costa et al., 2021). Other studies indicate that, in addition to the retreat and melting of sea ice in spring and summer, mostly driven by wind, light incidence and wind effects on the mixed layer also strongly influence phytoplankton bloom processes in the region (Ducklow et al., 2012, 2013).

Extreme weather events like the 2016 El Niño are expected to increase in both intensity and frequency due to climate change. Consequently, their impact on microbial communities is also likely to become more pronounced and recurrent. While the prokaryotic community appears to be more stable and less affected by environmental fluctuations, predictive models under IPCC warming scenarios for the Southern Ocean indicate significant changes. Fluctuations in *Nitrincolaceae* relative abundance, a bacterial family widely distributed in polar regions and recognized for its diverse carbon utilization strategies and rapid response to increased carbon availability, are expected (Thiele et al., 2022, 2023; Tonelli et al., 2021). This group was the only bacterial taxon consistently detected across all periods in this study, maintaining an average relative abundance above 2% (see Figure 4). Given its role as a keystone taxon in aquatic ecosystems, its decline could have important ecological consequences, potentially affecting carbon cycling and microbial community dynamics (Banerjee et al., 2018). Conversely, a rise in heterotrophic bacterial groups such as *Flavobacteriales* is anticipated (Tonelli et al., 2021). This taxon is closely associated with phytoplankton blooms and may have an advantage under warming conditions (Buchan et al., 2014; Landa et al., 2016; Tonelli et al., 2021).

### DIVERGING STABILITY IN PROKARYOTIC AND EUKARYOTIC COMMUNITIES OVER TIME

The eukaryotic fraction – represented by both Eukaryotic Phytoplankton (Chloroplast 16S) and Total Eukaryotic (Eukaryote 18S) – was the least stable and exhibited the greatest dissimilarity. In contrast, the prokaryotic fraction of the microbial community exhibited the greatest diversity stability over time, being dominated by the a few distinct ASVs belonging to the *Gammaproteobacteria* class (see Figures 2, 3 and 4). This trend for the prokaryotic community to exhibit greater stability compared to the eukaryotic community has also been observed in both subtropical and high-latitude regions (Signori et al., 2018; Y. C. Yeh & Fuhrman, 2022). The observed differences in temporal stability suggest that, overall, prokaryotic communities experience significantly stronger stabilizing influences compared to eukaryotic communities. This is likely due to the persistence of specific taxa, such as *SAR11* and *Nitrincolaceae*, which are consistently present across all samples and time periods with relatively high abundance (see Figure 4). These stabilizing factors may include larger population sizes, greater metabolic diversity, and higher dispersal potential relative to protists. Such characteristics likely confer resilience to prokaryotes, enabling them to survive or maintain small populations even in unfavorable environmental conditions (Leibold et al., 2004; Y. C. Yeh & Fuhrman, 2022).

### INTERRELATIONSHIPS BETWEEN PROKARYOTES AND EUKARYOTES

As previously noted, prokaryotic groups were found to be far more interconnected than eukaryotic groups, as observed in earlier studies (Luria et al., 2014, 2016). Intra-domain connections were less frequent, with interactions being either negative or positive. Negative interactions were observed exclusively between different domains (see Figure 5). Most of the negative relationships were observed between *Flavobacteriaceae*, *Rhodobacterales*, and eukaryotes—particularly photosynthetic ones. This may be linked to the fact that these prokaryotic groups are often associated with phytoplankton blooms, whether during periods of high or low abundance, depending on the bloom stage (Buchan et al., 2014; Landa et al., 2016). Furthermore, *Flavobacteriaceae* exhibited a strong negative correlation with parasite, saprotroph, and mixotroph groups, including *Pseudofungi*, *Picozoa* and *Dinophyceae*. These taxa appear to be signature groups associated with the late stages of phytoplankton blooms, when eukaryotic plankton biomass is being broken down by parasites and saprotrophs, which may explain their co-occurrence.

Interestingly, the *Acantharea* group showed approximately 20 positive interactions with various prokaryotic groups. However, *Acantharea* are also known for maintaining close symbiotic relationships with other eukaryotes, such as *Phaeocystis* (Brisbin et al., 2018). On the other hand, the bacterial group *Rickettsiales* showed only positive relationships with eukaryotes. *Rickettsiales*, an alphaproteobacterial order, are well known as obligate endosymbionts and parasites that infect a wide range of eukaryotic hosts (Schön et al., 2022).

Recently, researchers have argued that microbial interactions may play a more significant role in shaping community structure than environmental variables themselves (Gleich et al., 2025). These connections, whether positive or negative, underscore the importance of understanding not only the interactions themselves but also the broader consequences of environmental disturbances. An increase or decline in the relative abundance of a key taxon due to such disruptions can trigger cascading effects, reshaping the entire community structure and dynamics.

## MATERIAL AND METHODS

### SAMPLE COLLECTION

Within the maritime NAP, we sampled ten monitoring stations (M01-M10, Figure 6) across Drake South, Bransfield and Gerlach Straits. Our sampling regime covered various seasons. In 2013, samples were collected in February and November (summer and spring, respectively). In 2014, sampling took place in February, March, and November (summer, late summer, and spring, respectively). In 2015, samples were collected in February and March (summer and late summer, respectively). In 2016 and 2017, sampling occurred in February (summer), and in 2019, samples were collected in January (summer), resulting in a total of 105 samples (Supplementary Table 1). All samples were collected onboard the Brazilian Navy’s polar vessel Almirante Maximiano through the projects “Interactions in Marine Ecosystems near the Antarctic Peninsula under Different Impacts of Climate Change” (INTERBIOTA) and “Responses of the Pelagic Ecosystem to Climate Changes in the Southern Ocean” (EcoPelagos) as part of the Brazilian Antarctic Program. The map was generated with the ArcGIS version 10.4.1 program using Quantarctica version 3.2. (Matsuoka et al., 2021)

**Figure 6:**
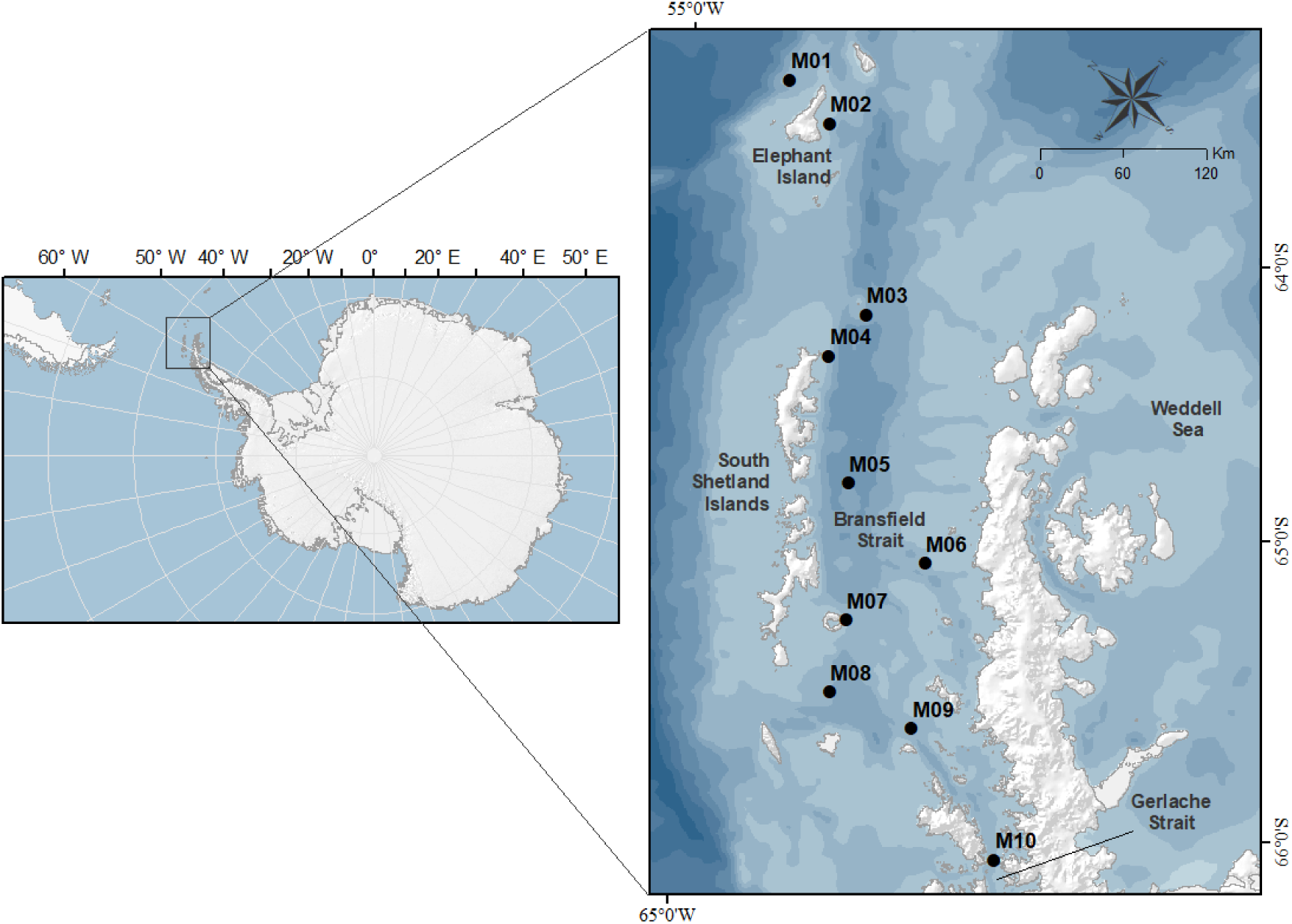
Sampling map illustrating the 10 monitoring stations (M01–M10). Samples were collected between 2013 and 2019. Lighter blue represents shallower areas, while darker blue indicates deeper regions. The deepest sampling station, M05, is approximately 1,525 m deep and the shallowest, M04, is approximately 160m deep.

At each sampling site, 3 L of surface seawater samples were collected using Niskin bottles from the top 5 meters. This was filtered through Sterivex^TM^ GP 0.2 μm filter unit (Millipore) using a peristaltic pump until the filters are completely dry, without any prior size fractionation or pre-filtration. The filters were immediately frozen onboard at -80°C until further laboratory analysis., A Sea-Bird CTD/Carousel 911 was deployed with each cast to collect data on temperature (°C) and salinity (PSU).

### DNA EXTRACTION

DNA from samples collected in the years 2013, 2014, and 2015 was extracted using the low-biomass protocol, with slight modifications (Boström et al., 2004; Manganelli et al., 2009; Signori et al., 2014). For the other years, the DNeasy PowerWater Kit (Qiagen, Hilden, Germany) was used following the manufacturer’s protocol.

### SEQUENCING METHODS AND PIPELINE

Relative abundance of microbial taxa was generated by PCR amplification of the V4 - V5 regions of the 16S rRNA and 18S rRNA genes from within the extracted DNA using universal primers 515F-Y (5′-GTGYCAGCMGCCGCGGTAA) and 926R (5′- CCGYCAATTYMTTTRAGTTT) (Parada et al., 2016). Samples were amplified using the following protocol (https://www.protocols.io/view/fuhrman-lab-515f-926r-16s-and18s-rrna-gene-sequen-vb7e2rn). In brief, PCR master mix included 9.5 µl of PCR water (VWR), 12.5 µl of GoTaq Master Mix (Taq DNA polymerase, 400μM dATP, 400μM dGTP, 400μM dCTP, 400μM dTTP and 3mM MgCl_2_), 2 µl of 1:1, 515F:926R barcoded primer mix (0.3 mM of each primer) and 1 µl of DNA per reaction for a final volume of 25 µl. PCR was performed under the following conditions: initial denaturing at 95°C for 120 seconds, followed by 30 cycles of 95°C for 45 seconds, 50°C for 45 seconds, and then 68°C for 90 seconds, and then a final elongation step at 68°C for 300 seconds. PCR products were then stored at 4°C before being cleaned using the Agencourt AMPure XP PCR purification protocol, and then DNA concentration was quantified using the Pico-Green dsDNA Quant-iT Assay Kit. Samples were then pooled and quantified using the Qubit dsDNA HS Assay Kit. Finally, the ratio of 16S:18S was calculated with the Bioanalyzer Chip: High Sensitivity DNA Kit.

Samples were sequenced using the Illumina HiSeq RapidRun2 x 250 technology at Tufts Medical School (http://tucf-genomics.tufts.edu/). After partial demultiplexing by the 6bp i7 index, sequences were fully demultiplexed with the in-line 5bp barcode using the following script (https://github.com/jcmcnch/demux-notes). Demultiplexed sequences were then denoised to amplicon sequence variants (ASVs) using a custom pipeline as described in (McNichol et al 2025). This pipeline is based on QIIME2 and DADA2 (Bolyen et al., 2019; Callahan et al., 2016) which can be found on GitHub (https://github.com/jcmcnch/eASV-pipeline-for-515Y-926R). We used a trim length of 220 for forward and 180 for reverse 16S and 18S reads so that ASVs could be directly compared to other datasets which use this pipeline, such as (McNichol et al., 2025). Finally, we employed two corrections to both 16S and 18S ASV tables before they were merged. The first adjusts for a known bias against longer 18S sequences during Illumina sequencing (Y. C. Yeh et al., 2021) and the second adjusts for random variations in sample quality using the DADA2 output statistics. For more details see (McNichol et al., 2025). Raw sequencing data is available on NCBI at BioProject ID: PRJNA1241329.

### DATA ANALYSIS

All statistical analysis was performed in R (4.4.0) (R Core Team, 2021). Samples were grouped by sampling areas (Drake South, Bransfield and Gerlache Straits) and/or sampling period (Late Summer, Spring, and Summer) and years (2013, 2014, 2015, 2016, 2017, 2018, 2019, Supplementary Table 1). To visualize the variation in salinity (PSU), and temperature (°C) across periods, boxplots were created using the *ggplot2* and *patchwork* packages within *tidyverse* (Wickham et al., 2019), as was the mean relative abundance of taxa comprising ≥2% of the total relative abundance for each period. To examine changes in alpha diversity within the Eukaryotic Phytoplankton (Chloroplast 16S), Total Eukaryotic (Eukaryote 18S), Prokaryotic (Prokaryote 16S), and MultiDomain (all Chloroplast 16S, Prokaryote 16S, and Eukaryote 18S) communities over time, Shannon H-indices were calculated based on relative abundance values and visualized with the *vegan* package and then plotted using *tidyverse* (Wickham et al., 2019).Finally, to investigate beta diversity and to investigate the influence of temperature on community structure, NMDS (non-Metric Multidimensional Scaling) analyses were performed using Bray–Curtis dissimilarity distances. The resulting plots included ellipses highlighting groups with >0.5 similarity, generated with the *tidyverse* (Wickham et al., 2019)*, vegan* (Oksanen et al., 2019), and *patchwork* packages (Pedersen, 2024). The *vegan* package was also used to perform a PERMANOVA to analyze which factors influenced the composition of the microbial community, using 999 permutations. We have also grouped the sequences into ecologically relevant plankton groups (McNichol et al., 2025) and determined Spearman’s R_S_ correlation coefficients using Mictools analysis (Albanese et al., 2018).

## CONCLUSION

This study highlights, for the first time, the dynamic nature of microbial communities in the maritime region of the Northwest Antarctic Peninsula, providing a multidomain perspective on its inter annual and seasonal variation. Our findings underscore the contrasting stability of prokaryotic and eukaryotic communities, with prokaryotes demonstrating remarkable resilience likely driven by intrinsic stabilizing factors such as metabolic diversity and dispersal potential. Conversely, the eukaryotic community, exhibited pronounced variability, reflecting their sensitivity to environmental shifts and seasonal changes. Despite the absence of significant differences in the measured environmental parameters, the observed temporal variations in community diversity and composition suggest the influence of unmeasured factors, such as wind-driven effects and nutrient availability. The dominance of specific taxa during bloom events, such as *Bacillariophyta* in 2016, highlights the influence of extreme weather events like the 2016 El Niño on microbial dynamics. Together, these findings provide critical insights, which can drive future studies into how Antarctic microbial communities may respond to future climate-driven disturbances, offering a foundation for understanding their ecological roles in a rapidly changing ocean.

## ACKNOWLEDGEMENTS

Thanks to the captains of Npo. Almirante Maximiano (H41) and their respective crews of the Brazilian Antarctic Expeditions “OPERANTAR XXXI, XXXII, XXXIII, XXXIV, XXXV and XXXIX”. We further thank the Laboratório de Estudos dos Oceanos e Clima (LEOC) at FURG for sharing their CTD data. This research was partially supported by the Projects INTERBIOTA (CNPq grant n° 407889/2013-2) and ECOPELAGOS (CNPq grant n° 442637/2018-7). LCF thanks the Scientific Committee on Antarctic Research (SCAR) for the Fellowship 2023-2024; the American Association of University Women (AAUW) for the International Fellowship 2022-2023 and the Coordenação de Aperfeiçoamento de Pessoal de Nível Superior – Brasil (CAPES) – Finance Code 001. JAF, YR, JM, and NLRW were supported by the Simons Collaboration on Computational Biogeochemical Modeling of Marine Ecosystems / CBIOMES) grant 549943 and the National Science Foundation (NSF) grant EF-2125142. CNS receives CNPq productivity fellowship.

---

**Supplementary Table 1:** Information on the sampling month, year, area, temperature, and salinity for each sample.

| SampleID | Area | Period | Temperature(C) | Salinity(PSU) |
| --- | --- | --- | --- | --- |
| M01-fev2014 | Drake South | 2014-Summer | 0.881 | 34.015 |
| M01-fev2015 | Drake South | 2015-Summer | 1.204 | 34.176 |
| M01-fev2017 | Drake South | 2017-Summer | -0.318 | 33.917 |
| M01-jan2019 | Drake South | 2019-Summer | 0.8885 | 34.3189 |
| M01-mar2014 | Drake South | 2014-LateSummer | 0.511 | 32.263 |
| M01-nov2013 | Drake South | 2013-Spring | -0.892 | 34.979 |
| M01-nov2014B | Drake South | 2014-Spring | -0.698 | 34.274 |
| M02-fev2014 | Drake South | 2014-Summer | 0.753 | 34.144 |
| M02-fev2015 | Drake South | 2015-Summer | 1.674 | 34.15 |
| M02-fev2016 | Drake South | 2016-Summer | 0.889 | 34.239 |
| M02-fev2017 | Drake South | 2017-Summer | 1.836 | 34.161 |
| M02-jan2019 | Drake South | 2019-Summer | 1.2644 | 34.3033 |
| M02-mar2014 | Drake South | 2014-LateSummer | 0.456 | 33.496 |
| M02-nov2013 | Drake South | 2013-Spring | -1.474 | 34.442 |
| M02-nov2014 | Drake South | 2014-Spring | -1.268 | 34.212 |
| M02-nov2014B | Drake South | 2014-Spring | -0.96 | 34.224 |
| M03-fev2014 | Bransfield | 2014-Summer | 0.311 | 33.746 |
| M03-fev2015 | Bransfield | 2015-Summer | 1.871 | 34.177 |
| M03-fev2016 | Bransfield | 2016-Summer | 0.729 | 34.123 |
| M03-fev2017 | Bransfield | 2017-Summer | 2.268 | 34.07 |
| M03-jan2019 | Bransfield | 2019-Summer | 0.9521 | 34.1112 |
| M03-mar2014 | Bransfield | 2014-LateSummer | 0.662 | 34.121 |
| M03-mar2015 | Bransfield | 2015-LateSummer | 1.064 | 34.022 |
| M03-nov2013 | Bransfield | 2013-Spring | -1.32 | 34.236 |
| M03-nov2013B | Bransfield | 2013-Spring | -1.251 | 34.301 |
| M03-nov2014 | Bransfield | 2014-Spring | -1.116 | 33.24 |
| M03-nov2014B | Bransfield | 2014-Spring | -0.791 | 34.168 |
| M04-fev2014 | Bransfield | 2014-Summer | 0.462 | 34.17 |
| M04-fev2015 | Bransfield | 2015-Summer | 1.374 | 33.904 |
| M04-fev2016 | Bransfield | 2016-Summer | 0.597 | 34.144 |
| M04-fev2017 | Bransfield | 2017-Summer | 1.734 | 34.103 |
| M04-jan2019 | Bransfield | 2019-Summer | 0.6875 | 34.244 |
| M04-jan2019A | Bransfield | 2019-Summer | 1.0816 | 34.1492 |
| M04-mar2014 | Bransfield | 2014-LateSummer | 0.948 | 33.847 |
| M04-mar2015 | Bransfield | 2015-LateSummer | 1.018 | 34.07 |
| M04-nov2013 | Bransfield | 2013-Spring | -1.235 | 34.084 |
| M04-nov2013B | Bransfield | 2013-Spring | -1.178 | 33.779 |
| M04-nov2014 | Bransfield | 2014-Spring | -1.012 | 36.336 |
| M04-nov2014B | Bransfield | 2014-Spring | -0.78 | 34.1 |
| M05-fev2014 | Bransfield | 2014-Summer | 1.085 | 33.976 |
| M05-fev2015 | Bransfield | 2015-Summer | 1.905 | 34.188 |
| M05-fev2016 | Bransfield | 2016-Summer | 0.553 | 34.295 |
| M05-fev2017 | Bransfield | 2017-Summer | 1.655 | 33.973 |
| M05-jan2019 | Bransfield | 2019-Summer | 1.1883 | 34.1102 |
| M05-mar2013 | Bransfield | 2013-LateSummer | 1.574 | 34.083 |
| M05-mar2014 | Bransfield | 2014-LateSummer | 0.59 | 33.89 |
| M05-mar2015 | Bransfield | 2015-LateSummer | 1.025 | 33.917 |
| M05-nov2013 | Bransfield | 2013-Spring | -1.282 | 34.112 |
| M05-nov2013B | Bransfield | 2013-Spring | -1.212 | 33.989 |
| M05-nov2014 | Bransfield | 2014-Spring | -0.93 | 34.861 |
| M05-nov2014B | Bransfield | 2014-Spring | -0.835 | 34.128 |
| M06-fev2014 | Bransfield | 2014-Summer | 0.185 | 33.92 |
| M06-fev2015 | Bransfield | 2015-Summer | 0.226 | 34.383 |
| M06-fev2016 | Bransfield | 2016-Summer | 0.216 | 34.385 |
| M06-fev2017 | Bransfield | 2017-Summer | 1.027 | 34.051 |
| M06-jan2019 | Bransfield | 2019-Summer | 0.946 | 34.3107 |
| M06-mar2014 | Bransfield | 2014-LateSummer | -0.573 |  |
| M06-mar2015 | Bransfield | 2015-LateSummer | 0.755 | 34.194 |
| M06-nov2013 | Bransfield | 2013-Spring | -1.596 | 34.145 |
| M06-nov2013B | Bransfield | 2013-Spring | -1.527 | 30.503 |
| M06-nov2014 | Bransfield | 2014-Spring | -0.987 | 34.387 |
| M06-nov2014B | Bransfield | 2014-Spring | -1.242 | 34.441 |
| M07-fev2014 | Bransfield | 2014-Summer | 0.812 | 33.905 |
| M07-fev2015 | Bransfield | 2015-Summer | 1.497 | 33.725 |
| M07-fev2016 | Bransfield | 2016-Summer |  |  |
| M07-fev2017 | Bransfield | 2017-Summer | 2.483 | 33.98 |
| M07-jan2019 | Bransfield | 2019-Summer | 1.0778 | 34.1473 |
| M07-mar2014 | Bransfield | 2014-LateSummer | 0.707 | 33.795 |
| M07-mar2015 | Bransfield | 2015-LateSummer | 1.02 | 34 |
| M07-nov2013 | Bransfield | 2013-Spring | -1.193 | 34.662 |
| M07-nov2013B | Bransfield | 2013-Spring | -1.772 | 34.549 |
| M07-nov2014 | Bransfield | 2014-Spring | -0.865 | 34.07 |
| M07-nov2014B | Bransfield | 2014-Spring | -0.823 | 34.092 |
| M08-fev2014 | Bransfield | 2014-Summer | 0.961 | 33.769 |
| M08-fev2015 | Bransfield | 2015-Summer | 1.605 | 33.849 |
| M08-fev2016 | Bransfield | 2016-Summer |  |  |
| M08-fev2017 | Bransfield | 2017-Summer | 2.392 | 34.07 |
| M08-jan2019 | Bransfield | 2019-Summer | 1.1887 | 34.0406 |
| M08-mar2014 | Bransfield | 2014-LateSummer | 0.749 | 33.228 |
| M08-mar2015 | Bransfield | 2015-LateSummer | 1.489 | 33.874 |
| M08-nov2013 | Bransfield | 2013-Spring | -1.326 | 34.036 |
| M08-nov2014 | Bransfield | 2014-Spring | -0.85 | 34.043 |
| M08-nov2014B | Bransfield | 2014-Spring | -1.07 | 34 |
| M08-nov201B3 | Bransfield | 2013-Spring | -1.317 |  |
| M09-fev2014 | Gerlache | 2014-Summer | 0.831 | 34.034 |
| M09-fev2015 | Gerlache | 2015-Summer | 0.727 | 34.264 |
| M09-fev2016 | Gerlache | 2016-Summer | 0.304 | 34.307 |
| M09-fev2017 | Gerlache | 2017-Summer | 1.599 | 34.158 |
| M09-jan2019 | Gerlache | 2019-Summer | 1.4225 | 34.0018 |
| M09-mar2014 | Gerlache | 2014-LateSummer | 0.423 | 33.319 |
| M09-mar2015 | Gerlache | 2015-LateSummer | 0.163 | 34.284 |
| M09-nov2013 | Gerlache | 2013-Spring | -1.205 | 34.104 |
| M09-nov2013B | Gerlache | 2013-Spring | -1.24 | 34.085 |
| M09-nov2014 | Gerlache | 2014-Spring | -0.686 | 34.031 |
| M09-nov2014B | Gerlache | 2014-Spring | -0.687 | 34.012 |
| M10-fev2014 | Gerlache | 2014-Summer | 1.005 | 33.617 |
| M10-fev2015 | Gerlache | 2015-Summer | 0.776 | 33.648 |
| M10-fev2016 | Gerlache | 2016-Summer | 0.593 | 33.94 |
| M10-fev2017 | Gerlache | 2017-Summer | 1.557 | 33.525 |
| M10-jan2019 | Gerlache | 2019-Summer | 1.2526 | 34.0537 |
| M10-mar2014 | Gerlache | 2014-LateSummer | 0.35 | 33.644 |
| M10-nov2013 | Gerlache | 2013-Spring | -0.881 | 34.029 |
| M10-nov2013B | Gerlache | 2013-Spring | -1.055 | 33.941 |
| M10-nov2014 | Gerlache | 2014-Spring | -0.474 | 34.29 |
| M10-nov2014B | Gerlache | 2014-Spring | -0.469 | 33.952 |

